# Regulation of yeast *RPL22B* splicing depends on intact pre-mRNA context of the intron

**DOI:** 10.1101/814301

**Authors:** Kateřina Abrhámová, Petr Folk

**Affiliations:** Department of Cell Biology, Faculty of Science, Charles University, Viničná 7, 128 00 Praha, Czech Republic

**Keywords:** intergenic regulation, pre-mRNA splicing, RNA structure, *Saccharomyces cerevisiae*, ribosomal protein genes

## Abstract

Yeast *RPL22A* and *RPL22B* genes form an intergenic regulatory loop modulating the ratio of paralogous transcripts in response to changing levels of proteins. Gabunilas and Chanfreau (Gabunilas and Chanfreau, PLoS Genet *12*, e1005999, 2016) and our group (Abrhámová et al., PLoS ONE *13*, e0190685, 2018) described that Rpl22 proteins bound to the divergent introns of *RPL22* paralogs and inhibited splicing in dosage dependent manner. Here, we continued to study the splicing regulation in more detail and designed constructs for *in vivo* analyses of splicing efficiency. We also tested Rpl22 binding to *RPL22B* intron in three-hybrid system. We were able to confirm the findings reported originally by Gabunilas and Chanfreau on the importance of a stem loop structure within the *RPL22B* intron. Mutations which lowered the stability of the structure abolished Rpl22-mediated inhibition. In contrast, we were not able to confirm the sequence specificity with respect to either Rpl22 binding or splicing inhibition within this region, which they reported. We contradict their results that the ‘RNA internal loop’ of *RPL22Bi* (nt 178CCCU181 and 221UGAA224) is crucial for mediating the Rpl22 effects. We assume that this discrepancy reflects the difference in constructs, as the reporters used by Gabunilas and Chanfreau lacked the alternative 5’ splice site as well as surrounding exons. Our own comparison confirms that deleting the sequence spanning alternative 5’ splice site lowers splicing efficiency, hinting to possible disturbances of the regulatory mechanism. We argue that the structural context of the ‘regulatory element’ may reach across the intron or into the surrounding sequences, similarly to what was found previously for other genes, such as *RPL30*. Apparently, more detailed analyses are needed to discern this intriguing example of splicing regulation.

## Introduction

Ribosomal proteins (RPs) constitute structural and regulatory components of ribosomes and as such remain highly conserved during evolution. Despite this constraint, some RPs acquired extra ribosomal functions by binding to RNAs other than rRNAs, mostly with impact on gene expression (Warner and McIntosh, 2009). The pre-mRNA molecules, their introns and untranslated regions in particular, can thus be viewed as evolving structures, which gained affinity for pre-existing RPs (Fahl et al., 2015). As it was found, Rpl26 bound p53 mRNA through its 5’ untranslated region (5’UTR), which stimulated translation and thus contributed to the regulation of DNA-damage response (Takagi et al., 2005). Rpl13a, upon IFN-γ activated phosphorylation, was released from ribosomes and translationally inhibited several groups of mRNAs as part of the GAIT system, which directed transcript selective translational control in myeloid cells (Arif et al., 2018; Mukhopadhyay et al., 2008).

While some RPs are unique, some RP coding genes (RPGs) underwent duplication and exist as paralog pairs. In yeast, which experienced whole genome duplication followed by loss of duplicates, RPGs, in their majority, retained their paralogs. These gene pairs were implicated in intergenic regulation (Parenteau et al., 2011). However, only a handful of such pairs were proven to regulate each other’s expression, such as *RPS14, RPS9*, and *RPL22* (Fewell and Woolford, 1999; Plocik and Guthrie, 2012; Petibon et al., 2016; Gabunilas and Chanfreau, 2016; Abrhámová et al., 2018). The incorporation of two different paralogs into ribosomes may give rise to ribosome heterogeneity, which became focus of recent attention. Proteomic and transcriptomic analyses found evidence for the existence of ribosomal subpopulations which differed in composition, interactomes, and perhaps mRNA specificity (Yadav et al., 2016; Shi et al., 2017; Segev and Gerst, 2018).

The propensity of RNA transcripts to form local structures of varying stability as well as interactions over long distances was studied for over 30 years (for reviews see Wan et al., 2011; Lin et al., 2016). More recently, structures begun to be examined in whole transcriptomes, using both chemical structure probing techniques as well as modelling approaches. Structures changed their parameters with cultivation conditions and upon stress, suggesting that RNA structural features might be of general significance for gene expression, including splicing and translation (Zhang et al., 2017; Rouskin et al., 2014; Kwok, 2016). Splicing sequences of 5’ splice site (5’ss), branch point (BP), or 3’ splice site (3’ss) can be incorporated into stems and thus blocked (Singh et al., 2006), or brought into proximity and made stronger (Lin et al., 2016; Rogic et al., 2008; Gahura et al., 2011). Exons can be looped out (Raker et al., 2009) or selected for alternative splicing through base-pairing interactions, such as in the case of multi-cluster mutually exclusive exons (Graveley, 2005; McManus and Graveley, 2011; Ivanov and Pervouchine, 2018).

*RPL22* paralogs represent an intriguing example of extra ribosomal function-gain and intergenic regulation throughout evolution. Extra ribosomal roles of Rpl22 paralogs were documented in yeast (see below), fruit fly, zebrafish, mice, and humans (Kearse et al., 2011; Mageeney et al., 2018; Zhang et al., 2017; O’Leary et al., 2013; Zhang et al., 2013). Human Rpl22 interacted with human telomerase RNA (Le et al., 2000) and was also sequestered, apparently in competition, by the Epstein-Barr virus (EBV)-encoded RNAs in EBV infected cells, possibly as part of the viral strategy to interfere with translation and growth (Fok, 2006; Houmani et al., 2009). Recently, Zhang and coworkers demonstrated that zebrafish Rpl22 and Rpl22-Like1 controlled morphogenesis during gastrulation. The proteins acted antagonistically to modulate splicing of Smad2 pre-mRNA, presumably in cooperation with HNRNP-A1 (Zhang et al., 2017).

In yeast, Rpl22A/B pair was shown to be part of oxidative stress response. Because *RPL22A* but NOT *RPL22B* mRNA is UUG rich, and because translation efficiency of UUG rich transcripts was increased under stress, the translation of *RPL22A* became more efficient, leading to change in A/B ratio (Chan et al., 2012). The effect was mediated by the methyl methanesulfonate (MMS) sensitive tRNA methyltransferase 9 (Trm9) (Begley et al., 2004, 2007). Rpl22 was also shown to be required for *IME1* mRNA translation and meiotic induction in *S. cerevisiae*, as *rpl22Δ* cells were unable to translate the *IME1* transcript because of its atypical 5’UTR (Kim and Strich, 2016). On the level of splicing regulated gene expression, the feedback-loop formed by the *RPL22A/B* pair coupled free [A]+[B] fluctuations to changes in [A]/[B]. The protein products of either paralog bound to *RPL22* introns, which caused moderate inhibition of the major paralog (*RPL22A*) and strong inhibition of the minor one (*RPL22B*) (Abrhámová et al., 2018). A/B changes were related to oxidative stress (see above), or disbalances in free RP concentration caused by exposures to MMS or Cd (Gabunilas and Chanfreau, 2016), supporting the interpretation that tuning the isoform ratio helps the cells to react to changing environment. What cellular process or machinery distinguishes the 19 amino acids difference between Rpl22A and Rpl22B remains unknown; the effects on splicing are apparently mediated by both A and B proteins (Abrhámová et al., 2018). Information on the interface between the intron and Rpl22, and between this complex and the spliceosome, is likewise limited. Gabunilas and Chanfreau postulated the intronic region between nt 153 and 225 (positions here and throughout the text are numbered relative to the first nucleotide of the *RPL22B* intron) as ‘regulatory element’ and found that it was responsible for protein binding (Fig3C, D in Gabunilas and Chanfreau, 2016). This is similar to our conclusions based on yeast three-hybrid assays (nt 165 to 236; see I2, Abrhámová et al., 2018). They also showed that a predicted stem forming between nt 182-188 and 214-220 is necessary for Rpl22-mediated regulation and can rescue splicing inhibition *in vitro* when added *in trans* as part of free RNA (nt 115-255) (Gabunilas and Chanfreau, 2016).

**Fig. 1.**
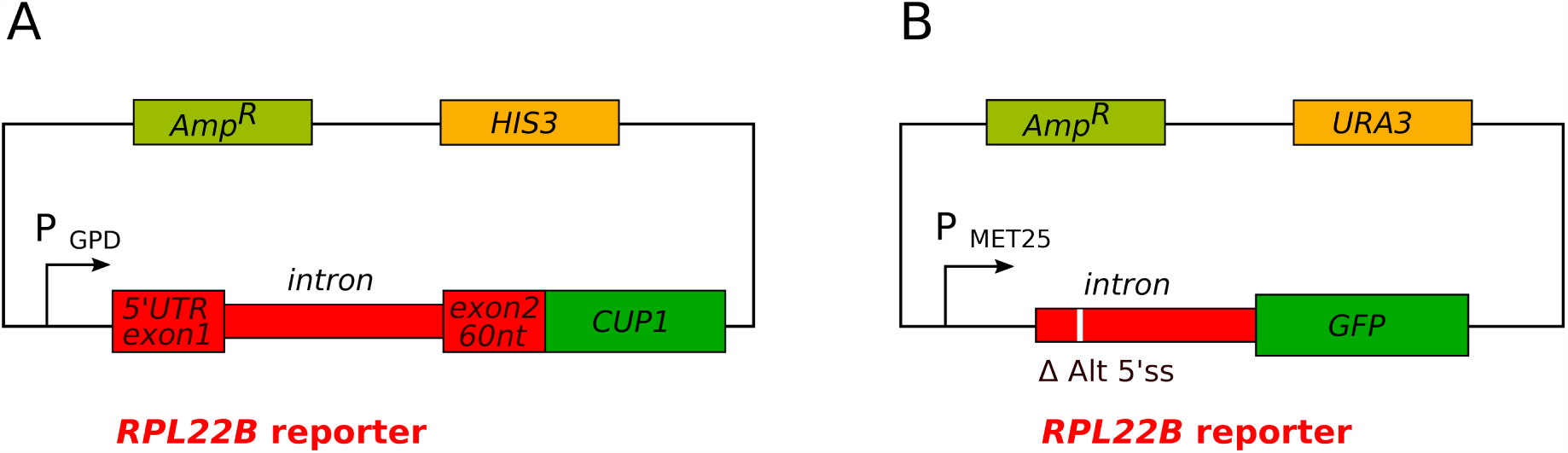
Comparison of reporter constructs used to analyze splicing of *RPL22B in vivo.* **(A)** In this study, we used splicing reporter constructs based on p423GPD backbone with *RPL22B* intron surrounded by its own 5’UTR, exon 1 and by 60 nts of exon 2, which were fused in frame with *CUP1* reporter (Abrhámová et al., 2018). Alternative 5’ss was preserved in all constructs unless indicated. **(B)** Gabunilas and Chanfreau used reporter plasmid derived from pUG23, where *RPL22B* intron without the surrounding sequences was inserted in front of the *GFP* reporter. In most of the constructs analyzed, alternative 5’ss was deleted (Gabunilas and Chanfreau, 2016).

**Fig. 2.**
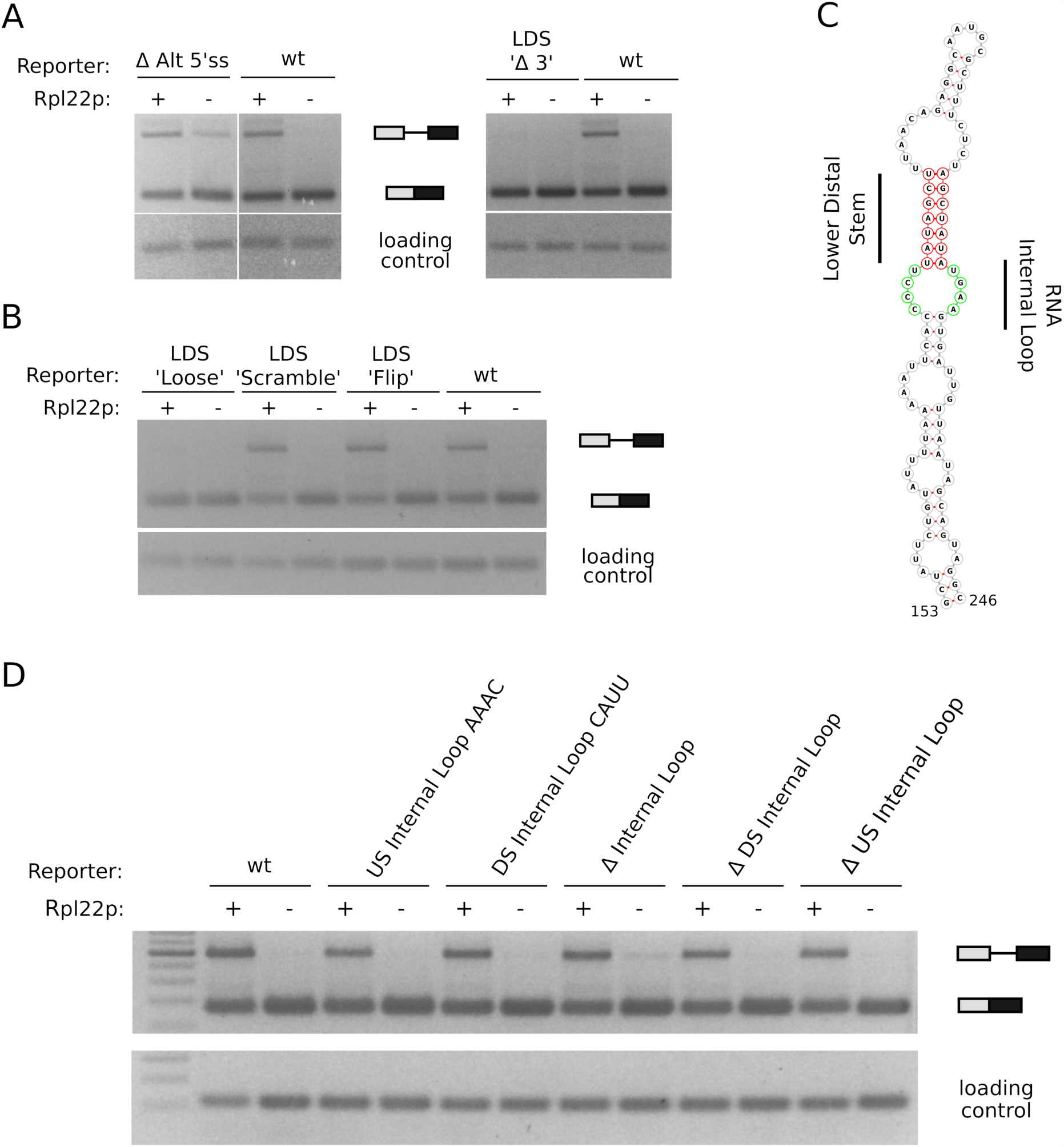
Splicing analysis of *RPL22B* reporter constructs. Splicing efficiency of *RPL22B-CUP1* reporters was tested in *rpl22a*Δ *rpl22b*Δ strain harboring pVTU260/RPL22A plasmid for Rpl22A overexpression and in the same strain transformed with empty vector. Semi-quantitative PCR was run on cDNA prepared using random hexamers from RNA isolated from exponentially growing cultures. *SPT15* and *SCR1* were used as loading control in A,B and D, respectively. One of at least three independent experiments is shown. **(A)** Alternative 5’ss is required for efficient splicing. Deletion of alternative 5’ss increased the level of unspliced RNA in the absence of Rpl22 as compared to WT (Δ Alt 5’ss). In comparison, deletion of 3’ strand of the ‘lower distal stem’ (LDS) maintained efficient splicing but almost completely canceled splicing inhibition by Rpl22 (LDS ‘Δ 3’). **(B)** Structure but not the sequence of LDS is important for splicing regulation. Mutants which maintained the pairing probabilities within the stem but changed its sequence (‘Scramble’, ‘Flip’) kept the regulation intact, but mutations that disrupted the structure of this part of the intron (‘Loose’ - mutation of nt 186GC to CG and 218AT to TA, ‘Δ 3’ - deletion of nt 214-220 in A) lost the regulation. **(C)** Predicted secondary structure of *RPL22Bi* in a region implicated in splicing regulation (nt 153 - 246). Structure was predicted by RNA fold (Lorenz et al., 2011) and visualized using Forna (Kerpedjiev et al., 2015). LDS and ‘RNA internal loop’ are labelled red and green, respectively. **(D)** In contradiction to previous findings (Gabunilas and Chanfreau, 2016), nucleotides within the ‘RNA internal loop’ are dispensable for *RPL22Bi* splicing regulation. CCCU to AAAC mutation of upstream part of the loop (‘US internal Loop AAAC’), UGAA to CAUU mutation in downstream part (‘DS Internal Loop CAUU’) as well as their deletions (deletion of both US CCCU and DS UGAA nucleotides – ‘Δ Internal Loop’, or *per partes* deletion of CCCU – ‘Δ US Internal Loop’ or UGAA – ‘Δ DS Internal Loop’) did not lose regulation upon Rpl22 protein overexpression.

**Fig. 3.**
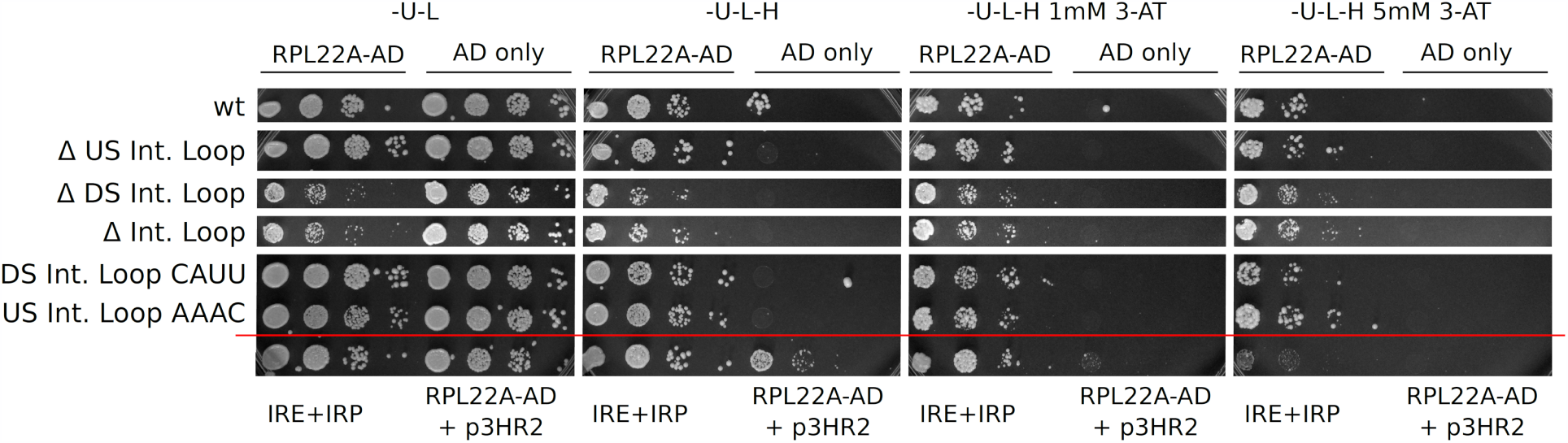
Yeast three-hybrid analysis of Rpl22 binding to *RPL22B* intron. Previously, we reported that Rpl22 binds the intron of *RPL22B* between nt 165 and 236 (Abrhámová et al., 2018). We introduced ‘RNA Internal Loop’ mutations in this region (see text and Fig2) and tested them in yeast three-hybrid system for their ability to interact with Rpl22. The system uses *HIS3* reporter gene, the activation of which depends on the recruitment of Rpl22 activation domain (AD) fusion protein through the interaction with the tested RNA. 10-fold serial dilutions of cells were spotted on plates with increasing concentrations of the metabolic retarder 3-aminotriazole (3-AT). ‘-U’, ‘-L’, and ‘-H’ denote the absence of uracil, leucine, and histidine in the medium. Cells with activation domain expressed alone (AD only, right parts of the panels) did not grow in the absence of histidine. Positive (IRE+IRP; SenGupta et al., 1996) and negative controls (Rpl22-AD with empty p3HR2 plasmid) are demarcated by red line (see also Abrhámová et al., 2018) for additional controls). Importantly, all RNA internal loop mutants supported cell growth to the same extent as the construct containing WT intron at 1 to 5 mM 3-AT.

Functional relationship of a ribosomal protein to both rRNA and its own intronic RNA, as found in *RPL22A/B*, is interesting for both structural and evolutionary biology. We began to characterize the intron-Rpl22 interaction in more detail, using constructs for *in vivo* measurements of splicing efficiency and yeast three-hybrid system. Our data agree with the findings of Gabunilas and Chanfreau (Gabunilas and Chanfreau, 2016) in that disrupting a stem structure within the *RPL22B* intron abolishes Rpl22-mediated inhibition. However, we contradict their conclusion that a sequence element within the ‘RNA internal loop’ (*RPL22Bi* nt 178CCCU181 and 221UGAA224) is crucial for Rpl22 binding and splicing inhibition. We confirmed that the intron’s response is dependent on the presence of intact alternative 5’ splice site (ALT5’ss), which suggests that the inhibitory function of the intron involves not only the 182-188/214-220 stem, but also the 5’ proximal region of the intron and potentially upstream parts of the transcript.

## Results

We decided to study the properties of the *RPL22* introns in more detail, using sensitive *in vivo* reporter assay and yeast three-hybrid analysis. Our aim is to decipher the mechanism which *RPL22* derived RNA molecules use in the regulation of expression of their own genes.

We set out to design splicing constructs harboring *RPL22B* intron surrounded by its own 5’UTR and exon 1 and by 60 nt of exon 2 fused in frame with *CUP1* reporter on the intron’s 5’ and 3’ ends, respectively (see Fig1A). In this way, any structural and functional features of the pre-mRNA molecule between nt −56 and +381 (positions are numbered relative to the first nucleotide of the *RPL22* intron) were preserved, including ALT5’ss (65-GTTTGT). We introduced mutations in regions implicated in Rpl22 mediated regulation and binding and assessed their impact on splicing. We employed cells with both endogenous *RPL22* paralogs deleted, which were harboring an expression plasmid, either empty or bearing *RPL22A*. We could thus test the splicing of *RPL22B* in the absence or the presence of Rpl22A and directly compare the readouts. This setup allowed us to differentiate mutations which were (i) splicing inhibitory regardless of the presence or absence of Rpl22 protein from those which were (ii) splicing permissive in its presence – rendering the intron unresponsive to Rpl22-mediated inhibition. In addition, some mutations (iii) showed graded effect and some (iv) affected the relative usage of alternative splice site.

One example of splicing permissive mutation is the destabilization of a hypothetical stem between nt 182-188 and 214-220 of *RPL22Bi* (Fig2A, B). This stem can be predicted using both RNA fold and Mfold (Lorenz et al., 2011; Zuker, 2003). To facilitate comparison between the work of Gabunilas and Chanfreau and our own when referring to mutations, we adopted the denotations used by these authors (Fig2C). We found that deleting the nucleotides 214-220 of the 3’ arm of the ‘lower distal stem’ (LDS) turned the intron refractory to Rpl22A inhibition. To further confirm that LDS is important for the negative regulation of *RPL22B* intron splicing, we prepared constructs coding for stems of similar predicted stability but with the LDS arms flipped (LDS’Flip’) or scrambled (LDS’Scramble’) to eliminate any nucleotide-specific effects. As shown in Fig2B, these constructs retained WT behavior. In contrast, mutations leading to decreased pairing probability within LDS (LDS ‘Δ 3’, LDS ‘Loose’) led to splicing permissive outcome – the intron’s responsiveness to Rpl22A was almost lost.

Targeting the ALT5’ss (deletion of nt 65-70) impaired splicing (Fig2A). We observed the accumulation of unspliced pre-mRNA even in the absence of the inhibitory protein in comparison with WT construct, where pre-mRNA was not detected. The mutant construct, however, still maintained some regulatory potential as the unspliced pre-mRNA signal was more pronounced in the presence of Rpl22A. The result showing that the deletion of ALT5’ss negatively affects splicing efficiency of *RPL22B*i was already obtained by Chanfreau group (Fig5B in Kawashima et al., 2014; FigS2C in Gabunilas and Chanfreau, 2016).

In addition to LDS, Gabunilas and Chanfreau targeted several other parts of the structured region between nt 153 and 225 of *RPL22Bi*. Based on their results, the authors proposed a more detailed model of *RPL22Bi*-Rpl22A interaction (FigS4 and S5 in Gabunilas and Chanfreau, 2016). We were concerned by the fact that their constructs did not include the neighboring regions of the intron and, most importantly, contained the deletion of ALT5’ss, shown by both groups to change splicing of *RPL22Bi*. We therefore decided to implement the above mentioned modifications in our constructs and test their behavior under our conditions, using the entire *RPL22B* intron with ALT5’ss and the surrounding sequences. We prepared constructs DS Internal Loop CAUU (UGAA to CAUU mutation), US Internal Loop AAAC (CCCU to AAAC mutation), Δ Internal Loop (deletion of the entire ‘RNA internal loop’) as well as Δ DS Internal Loop and Δ US Internal Loop, where the 5’ or 3’ arm of the ‘RNA internal loop’ is deleted. We found that none of the mutations disrupted negative splicing regulation by Rpl22A because all the reporters reacted to the Rpl22A overexpression by increasing the amount of unspliced pre-mRNA to levels obtained in the WT reporter (Fig2B). In the uninhibited state, they were also spliced with the same efficiency as the WT intron; possible exception was Δ Internal Loop mutant, which showed slightly less efficient splicing.

Rpl22A binds to parts of *RPL22i*, as was shown by Gabunilas and Chanfreau as well as by us. While there is still no ultimate proof that the binding is direct and not mediated or aided by other molecules, it can be reproduced in yeast three-hybrid system using only fragments of the intron. Rpl22A binding is dependent on the presence of lysines at the protein’s RNA-binding interface, as mutants where these lysines are substituted by glutamates are binding-negative in the three-hybrid system and unable to inhibit *RPL22B* splicing *in vivo* (Abrhámová et al, 2018). We used three-hybrid system to test Rpl22A binding to the mutant versions of *RPL22Bi* (Fig3). None of the mutations targeting the ‘RNA internal loop’ (Fig2D) impaired the ability of this part of intron to activate reporter gene synthesis in yeast three-hybrid system, implying that they were still compatible with Rpl22 binding.

## Discussion

Important aspect of *RPL22A/B* biology is the differentially regulated splicing of their pre-mRNAs. *RPL22* introns evolved into regulatory elements which block pre-mRNA maturation at high Rpl22 concentration. We would like to understand the mechanism of inhibition as it may be of general significance for the splicing field.

We constructed splicing reporters for *in vivo* testing of mutant introns and begun to characterize the elements which are necessary and sufficient for the Rpl22-dependent regulation (Fig1A). We found that deletion of the ALT5’ss of *RPL22Bi* impairs splicing efficiency at the major site (Fig2A). In previously published studies, we and others have shown that splicing *in vivo* uses alternative 5’ as well as 3’ splice sites, albeit at low frequencies (Abrhámová et al., 2018; Kawashima et al., 2014; Schreiber et al., 2015; Gould et al., 2016; Aslanzadeh et al., 2018). While the alternative site does not lead to a functional product, it may be part of the spliceosome assembly process, e.g., as an adjunct assembly landing platform (Libri et al., 2000; Spingola and Ares, 2000). In all our constructs, we retain the ALT5’ss and also include intron proximal sequences.

The inclusion of intron-surrounding sequences and the presence of intact ALT5’ss represent important differences in approach between Gabunilas and Chanfreau and our lab. On the beginning of their search for the region responsible for the regulation of *RPL22Bi* splicing, Gabunilas and Chanfreau prepared deletions lacking the first (Δ7-152) or the second (Δ153-297) half of the sequence between 5’ss and BP. The construct Δ7-152 retained the capacity to inhibit splicing in the presence of *RPL22A*, but it caused increased accumulation of unspliced transcript in *rpl22a*Δ cells. The construct Δ153-297, on the other hand, gave less clear result and showed increased usage of ALT5’ss (FigS2A in Gabunilas and Chanfreau, 2016). At this point, the authors decided to delete the ALT5’ss (FigS2C in Gabunilas and Chanfreau, 2016). Their own results show, however, that ΔALT5’ss-intron is splicing-impaired, both in the presence and absence of Rpl22A (compare lanes 1 to 3 and 2 to 4 in FigS2C in Gabunilas and Chanfreau, 2016). The Δ153-297ΔALT5’ss construct showed relieved inhibition in WT cells, but unspliced pre-mRNA continued to be increased in the absence of Rpl22A. Strikingly, the deletion of ALT5’ss had opposite impact on splicing efficiency of WT intron (inhibition of splicing in both WT and *rpl22aΔ* cells) in comparison with its truncated version (Δ153-297; improvement of splicing). We believe that their as well as our deletion constructs show that the region between 5’ss and BP, including the ALT5’ss, is required for full regulatory capacity of the intron. Notably, mutations or ablations of intron-adjacent exon sequences were shown to impact splicing efficiency (Crotti et al., 2007; Chanfreau et al., 1999). It was also found that splicing of *RPS9* was regulated by the interplay between an intronic structure and 3’ untranslated region (Petibon et al., 2016). The results on *RPL22Bi*, where both ALT5’ss deletion and other manipulations are present simultaneously, may thus be difficult to interpret unless we understand the structure-function relationships in detail.

We assume that the above differences may be the reason why our results on the mutations of the ‘RNA internal loop’ contradict the conclusions of Gabunilas and Chanfreau. We found that the constructs US Internal Loop AAAC and DS Internal Loop CAUU both maintained their regulatory potential (Fig2D), as did Δ DS Internal Loop and Δ US Internal Loop. Only Δ Internal Loop construct showed slight decrease of splicing efficiency in the absence of Rpl22, but even this construct inhibited splicing in response to Rpl22 overexpression. Gabunilas and Chanfreau first tested Δ191-211 UUCG ΔInternal Loop mutant, which was in their assay more splicing permissive than Δ191-211 UUCG mutant, suggesting to them that the internal loop nucleotides are important for splicing regulation. However, these constructs lacked the apical portion of the stem between nt 191 and 211, which obviously made them fold differently from the full-length stem, irrespective of the internal loop deletion. Second, they found that DS Internal Loop CAUU manipulation is more splicing permissive than US Internal Loop AAAC (FigS5B in Gabunilas and Chanfreau, 2016), which led them to propose that the sequence GUAA (mutated in CAUU) is crucial for mediating the effect of Rpl22A. Unfortunately, Gabunilas and Chanfreau did not include comparisons between WT, *rpl22aΔ* and Rpl22A overexpressing cells (Fig3, FigS4 and S5 in Gabunilas and Chanfreau, 2016). This makes it difficult to judge on the capacity of the constructs to regulate in response to changing Rpl22A concentration.

To complement our findings in Fig3C, we tested all our mutants (see also above) for their capacity to bind Rpl22 in three-hybrid assay. Indeed, none of the mutations hampered Rpl22A binding; the binding strength, based on the ability of reporter cells to grown on 1-5mM 3-AT, was identical to WT. This further supports our assumption that the mutations of the ‘RNA internal loop’ nucleotides, including its DS arm GUAA (nt 178CCCU181 and 221UGAA224), do not disturb Rpl22A mediated inhibition.

The regulatory mechanism of the intron may encompass not only the stem between nt 182-188 and 214-220 but also 5’ upstream parts of the intron including the ALT5’ss. Splicing was shown to be modulated from regions outside of the introns themselves through (i) SR and other proteins binding outside of introns to exonic splicing silencers/enhancers (for review see Lee and Rio, 2015), and (ii) RNA structures involving but not limited to exons (AbuQattam et al., 2018). In one of the well documented examples from *S. cerevisiae*, ribosomal Rpl30 protein binds to a stem-loop structure formed within the first exon and the 5’ end of the intron of *RPL30* transcript. The binding of the protein was shown to modulate both splicing (Eng and Warner, 1991; Vilardell and Warner, 1994) and translation, as the stem-loop can apparently form even after splicing (Eng and Warner, 1991; Mao, 1999; Vilardell and Warner, 1994; Vilardell et al., 2000). The effects of Rpl30 on *RPL30* splicing, which are brought about by RNA structure involving both intron and exon, can be taken as an illustration that the mechanism regulating splicing can be understood only by taking into account the context of the whole gene.

The results presented in this communication illustrate the requirement of ALT5’ss for *RPL22Bi* regulation. At the same time, they do not support the role of ‘RNA internal loop’ as the crucial interface required for Rpl22-triggered mechanism. Our results agree with Gabunilas and Chanfreau on the role of the ‘lower distal stem’, despite the differences in constructs. We assume that the structural disturbance (of making the sequences of LDS pairing-incompatible) is so substantial that it melts the RNA structure, eliminating both Rpl22A binding as well as any potential inhibitory effects. The involvement of ALT5’ss for *RPL22Bi* regulation would also explain the failure of Gabunilas and Chanfreau to show that their ‘regulatory element’ can function in an unrelated intron (*RPS21A*; Gabunilas and Chanfreau, 2016).

## Methods

### Yeast strains and cultivation methods

Yeast strains were transformed using lithium acetate method (Gietz and Schiestl, 2007). Plasmids used in this study are listed in Table 1. *rpl22aΔ rpl22bΔ* strain was prepared as described previously (Abrhámová et al., 2018). For splicing analysis, yeast cells were inoculated from overnight grown pre-cultures and let to grow for two generations in synthetic medium without uracil and histidine. Yeast three-hybrid system testing was done essentially as described previously (Abrhámová et al., 2018).

**Table 1.**
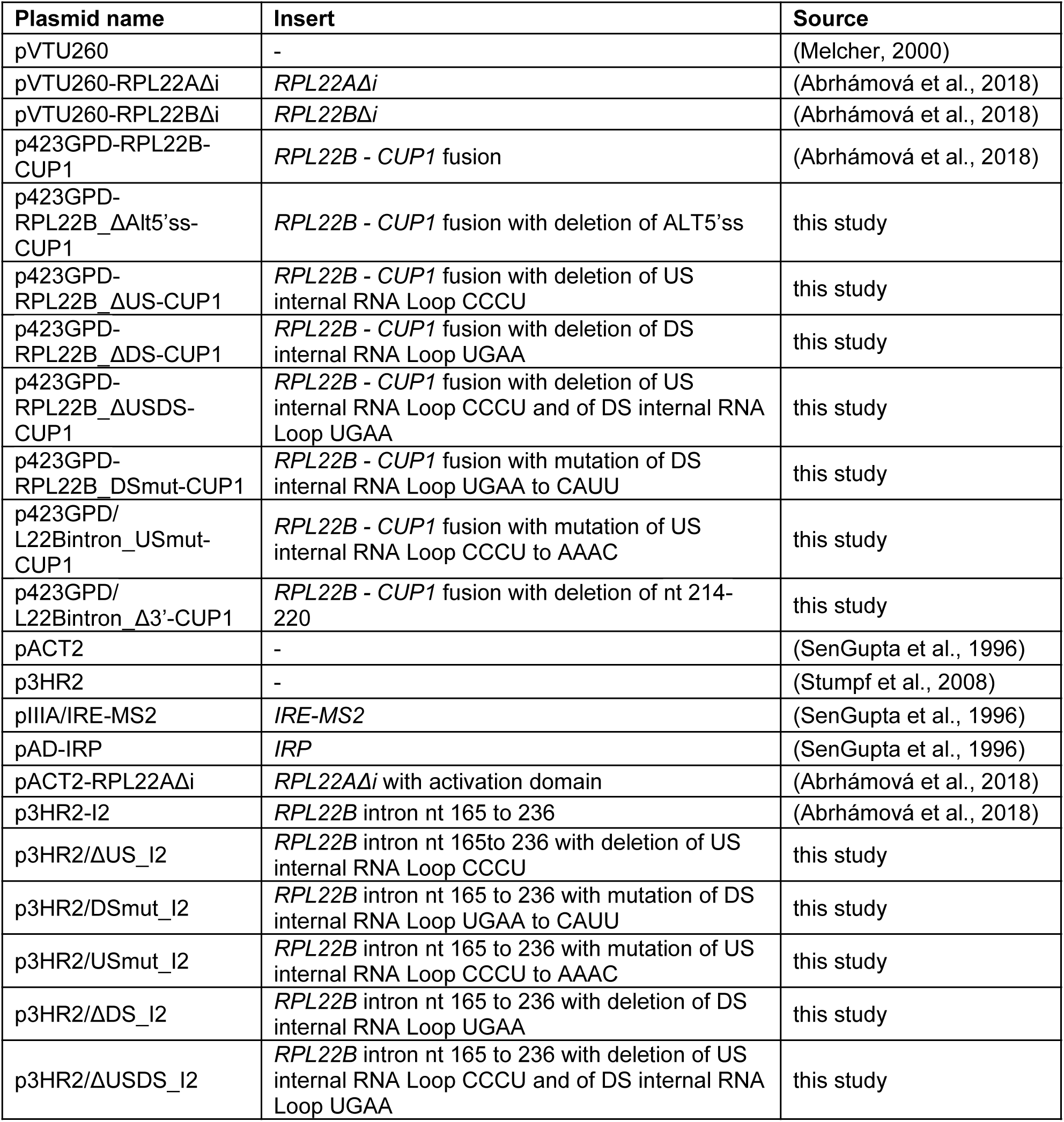
List of plasmids used in this study.

| **Plasmid name** | **Insert** | **Source** |
| --- | --- | --- |
| pVTU260 | - | (Melcher, 2000) |
| pVTU260-RPL22AΔi | *RPL22AΔi* | (Abrhámová et al., 2018) |
| pVTU260-RPL22BΔi | *RPL22BΔi* | (Abrhámová et al., 2018) |
| p423GPD-RPL22B-CUP1 | *RPL22B - CUP1* fusion | (Abrhámová et al., 2018) |
| p423GPD-RPL22B_ΔAlt5'ss-CUP1 | *RPL22B - CUP1* fusion with deletion of ALT5'ss | this study |
| p423GPD-RPL22B_ΔUS-CUP1 | *RPL22B - CUP1* fusion with deletion of US internal RNA Loop CCCU | this study |
| p423GPD-RPL22B_ΔDS-CUP1 | *RPL22B - CUP1* fusion with deletion of DS internal RNA Loop UGAA | this study |
| p423GPD-RPL22B_ΔUSDS-CUP1 | *RPL22B - CUP1* fusion with deletion of US internal RNA Loop CCCU and of DS internal RNA Loop UGAA | this study |
| p423GPD-RPL22B_DSmut-CUP1 | *RPL22B - CUP1* fusion with mutation of DS internal RNA Loop UGAA to CAUU | this study |
| p423GPD/L22Bintron_USmut-CUP1 | *RPL22B - CUP1* fusion with mutation of US internal RNA Loop CCCU to AAAC | this study |
| p423GPD/L22Bintron_Δ3'-CUP1 | *RPL22B - CUP1* fusion with deletion of nt 214-220 | this study |
| pACT2 | - | (SenGupta et al., 1996) |
| p3HR2 | - | (Stumpf et al., 2008) |
| pIIIA/IRE-MS2 | *IRE-MS2* | (SenGupta et al., 1996) |
| pAD-IRP | *IRP* | (SenGupta et al., 1996) |
| pACT2-RPL22AΔi | *RPL22AΔi* with activation domain | (Abrhámová et al., 2018) |
| p3HR2-I2 | *RPL22B* intron nt 165 to 236 | (Abrhámová et al., 2018) |
| p3HR2/ΔUS_I2 | *RPL22B* intron nt 165 to 236 with deletion of US internal RNA Loop CCCU | this study |
| p3HR2/DSmut_I2 | *RPL22B* intron nt 165 to 236 with mutation of DS internal RNA Loop UGAA to CAUU | this study |
| p3HR2/USmut_I2 | *RPL22B* intron nt 165 to 236 with mutation of US internal RNA Loop CCCU to AAAC | this study |
| p3HR2/ΔDS_I2 | *RPL22B* intron nt 165 to 236 with deletion of DS internal RNA Loop UGAA | this study |
| p3HR2/ΔUSDS_I2 | *RPL22B* intron nt 165 to 236 with deletion of US internal RNA Loop CCCU and of DS internal RNA Loop UGAA | this study |

### Splicing analyses

RNA isolation and reverse transcription were performed as described previously (Abrhámová et al., 2018). Semiquantitative PCR was run in 25μl-reactions with 5μl of 50 times diluted cDNA as template and with primers listed in Table2 for 25 to 28 cycles. The whole sample was loaded on 2.5% agarose gel and DNA was stained by ethidium bromide. Pictures were taken by Panasonic DMC-FZ7 camera, processed in GNU Image Manipulation Program 2.10.6 (https://www.gimp.org) and assembled in Inkscape 0.92.3 (https://www.inkscape.org).

### Plasmid preparation

*RPL22B* intron mutants were synthetized by GeneArt (Thermo Fisher Scientific) and swapped using BamHI and EcoRI restriction sites with WT version of *RPL22B* in p423GPD-RPL22B-CUP1. For three-hybrid testing, mutated versions of *RPL22B* were amplified from GeneArt DNA Strings with primers specific for each mutant (listed in Table 2) and cloned in SphI site in p3HR2 vector. Cloning outcome was verified by restriction analysis and sequencing.

**Table 2.**
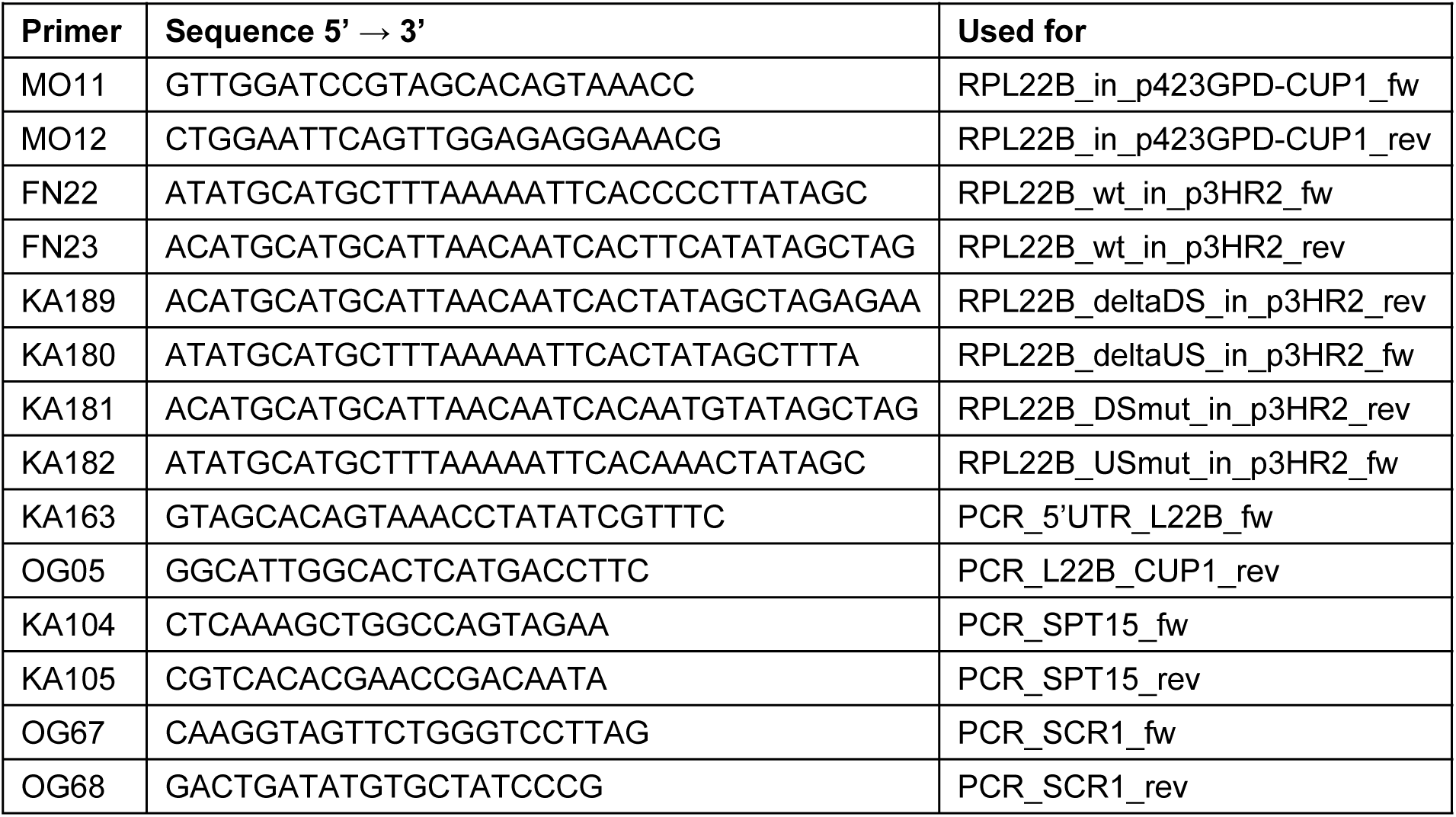
List of primers used in this study.

## Acknowledgments

We are grateful to Eva Krellerová for technical assistance. This work was supported by Charles University PROGRES program Q43 and UNCE 204013.

